# Transcriptional changes in human palate and skin healing

**DOI:** 10.1101/2022.09.01.506279

**Authors:** Trevor R Leonardo, Lin Chen, Megan E Schrementi, Junhe Shi, Phillip T Marucha, Kimberly Glass, Luisa A DiPietro

## Abstract

Most human tissue injuries lead to the formation of a fibrous scar and result in the loss of functional tissue. One adult tissue that exhibits a more regenerative response to injury with minimal scarring is the oral mucosa. We generated a microarray gene expression dataset to examine the response to injury from human palate and skin excisional biopsies spanning the first seven days after wounding. Differential expression analyses were performed in each tissue to identify genes overexpressed or underexpressed over time compared to baseline unwounded tissue. To attribute biological processes of interest to these gene expression changes, gene set enrichment analysis was used to identify core gene sets that are enriched over the time-course of the wound healing process with respect to unwounded tissue. This analysis identified gene sets uniquely enriched in either palate or skin wounds and gene sets that are enriched in both tissues in at least one time point after injury. Finally, a cell-deconvolution analysis was performed to better understand the cell type distribution in these tissues and how it changes over the time course of wound healing. This work provides a source of human wound gene expression data that includes two tissue types with distinct regenerative and scarring phenotypes.

## Introduction

A fully regenerative response to injury remains an unattainable outcome in most animals. The observation that most tissues in adult mammals heal with a scar is well established. One adult tissue that exhibits enhanced wound healing is the oral mucosa. Previous studies in human, murine, and pig models of wound healing have demonstrated that the oral mucosa exhibits a more rapid wound closure, less angiogenesis, decreased inflammation, and reduced scarring when compared to skin (1-8). Since the oral mucosa and skin reside in distinctly different environments, tissue milieu has been suggested to be responsible for the differences in wound healing at the two sites. However, skin transposed into the oral cavity maintains its morphologic characteristics (9), and may result in intraoral keloids (10), suggesting that rapid wound healing in the oral mucosa is not derived simply from the milieu. Intrinsic differences in the injury response of isolated cells from skin and mucosa have been demonstrated in multiple studies (1, 2, 11). For example, compared to skin, isolated oral mucosal epithelial cells show increased migration and proliferation (5), and isolated fibroblasts from the two sites exhibit different contractile properties (12).

To fully capture and compare the changes that occur after injury in human oral and skin tissues, a temporal analysis of the transcriptome would be necessary. However, few studies have produced human transcriptomic datasets that include multiple time points over the course of the wound healing process (3, 6, 13). Our lab has previously shown in a murine model that the temporal transcriptomic response in tongue and skin wound healing is intrinsically different (4). Recent studies of human tissues show similar differences between oral and skin wounds (6). This prior study identified Paired like homeodomain 1 (PITX1) and SRY-box transcription factor 2 (SOX2) as transcription factors responsible for priming the oral mucosal epithelium for wound repair due to their high expression levels in oral mucosa at baseline compared to skin and activation of downstream signaling pathways (14). Other studies have identified a specific fibroblast lineage designated by the expression of transcription factor engrailed-1 (EN1) as a key driver in the scarring outcomes in skin (15). Despite these and other significant advances in knowledge of oral healing, the problem of scarring remains unsolved and is a costly burden on society (16, 17).

We hypothesized that the transcriptional changes in the temporal response to injury in human palate and skin are distinct, and that different gene sets are enriched in a time- and tissue-specific manner. In order to gain a better understanding of the temporal changes of such a complex set of processes, we generated and analyzed a wound healing microarray gene expression dataset from human palate and skin excisional biopsies that spans the first seven days of healing. This analysis compares the transcriptomic response to injury between palate and skin tissues and investigates the functional changes that occur at the gene set level in a palate-specific, skin-specific, or shared manner. Finally, a cell-deconvolution analysis was performed to infer temporal changes in cell type distribution in palate and skin healing.

## Materials and methods

### 1. Study design and tissue collection

All procedures were carried out at the Center for Wound Healing and Tissue Regeneration in the Department of Periodontics at the University of Illinois Chicago (UIC), and had ethical approval by the university institutional review board (IRB). Volunteer participants were recruited from the student, staff, and patient population at UIC. Participants between 18-35 years of age were screened using a survey-form, which contained information regarding demographics and general health. Individuals were ineligible for the study if they: currently used tobacco products, had an oral disease needing emergency treatment, had a known skin disease other than acne, or had a medical problem that made them a high surgical risk. Furthermore, females were ineligible for the study if they were currently pregnant, trying to become pregnant, or breast-feeding. All wounding procedures were performed by a periodontist. Wounds were placed between 10-11 a.m. to minimize variability. Individuals were asked to refrain from alcohol and non-prescription drugs for a period of 24 hours prior to the experimental sessions. One skin and one mucosal wound were placed on each subject. For oral wounds, the side of the mouth was chosen by coin toss. The wounding site was anesthetized using 2% lidocaine. A mucosal wound measuring 1×5 millimeters (mm) was placed between the first and second molars, approximately 3 mm from the marginal gingiva using a Bard Parker handle, with two blades affixed 1 mm apart to assure uniformity of the wounds. A single blade scalpel was then used to make a relatively uniform 1.5 mm deep wound, removing the biopsy tissue. The wounds were not dressed, and subjects were instructed not to change their normal oral hygiene procedures, with the exception of avoiding the use of alcohol-based mouthwash. A 2×5 mm longitudinal excisional biopsy was made using a 2 mm two blade scalpel to collect oral wound biopsies. The dermal wound site was on the ventral side of the non-preferred forearm, approximately 4 centimeters below the elbow. The biopsy site was anesthetized with 2% lidocaine. Dermal wounds were created in a similar fashion to mucosal wounds, a 1×5 mm longitudinal wound approximately 1.5 mm deep. Wounds were dressed for one day with a standard adhesive bandage. Dermal and oral wounds were biopsied at 6 hours, Day 1, Day 3, and Day 7 post-wounding. Unwounded tissue (time 0) was also processed. A separate wound was used for each biopsy.

### 2. RNA isolation, microarray data collection and annotation

Tissue biopsies were collected in RNAlater solubilizing reagent according to manufacturer’s instructions (Thermo Fisher Scientific, Waltham, MA). Total cellular RNA was isolated from collected samples using RNeasy column-based kits (Qiagen, Hilden, Germany) according to manufacturer’s instructions and stored at -80°C until all samples were collected. Samples were analyzed for RNA quantity and quality using the Experion system (Bio-Rad, Hercules, CA) per manufacturer’s instruction. Transcriptional profiling was performed with the use of the Affymetrix Human Genome U133 Plus 2.0 GeneChip array (Affymetrix, Santa Clara, CA). All labeling reactions and hybridizations were carried out according to the standard GeneChip® eukaryotic target labeling protocol. Quality control of the 105 raw (.CEL) data files was performed using the web-based application ArrayAnalysis.org (18) which led to exclusion of 9 samples, retaining 96 samples for further analysis. Probes were mapped to the custom chip description file (CDF) HGU133Plus2_Hs_ENSG (Version V22.0.0) from brainarray (19), followed by normalization using the robust multichip averaging (RMA) procedure in the R/Bioconductor affy (20) package. Genes without annotated gene symbols were removed from the analysis, leaving a total of 18,117 annotated genes.

### 3. Differential expression and functional enrichment analysis

The R/Bioconductor software package linear models for microarray data (LIMMA) (21, 22) was used to conduct all differential expression comparisons on the normalized microarray data. Subjects were treated as a random effect using the duplicateCorrelation function and input into the linear model fit. Individual contrasts were designed to compare the unwounded palate and skin groups, along with each individual time point after injury versus their respective unwounded tissue group. This design allowed us to correct for baseline tissue-specific gene expression values and accurately extract the transcriptomic wound response in palate and skin. Multiple hypothesis testing across genes was corrected for separately in each contrast using the Benjamini & Hochberg (BH) method (23) to control for false discovery rate (FDR). An adjusted p-value ≤ 0.05 was used as the threshold for statistical significance. Genes with an absolute log2 fold change (log2FC) ≥ 1 were used for K-means clustering and heatmap generation. A cluster number of k=5 was determined to be suitable based on results of the Elbow method. The t-statistics generated from each contrast were ranked in descending order for all genes and input into the fast Gene Set Enrichment Analysis (fGSEA) (24) R package using the pre-ranked GSEA function fgseaMultilevel with default settings. Gene sets included were the Hallmark gene set collection (v7.0), the Gene Ontology (GO) biological processes (v7.0) and the curated Kyoto encyclopedia of genomes and genes (KEGG) gene sets (v7.0) downloaded from the Molecular Signatures Database (MSigDB) on March 4^th^ 2020 (25, 26). A BH adjusted p-value of ≤ 0.05 was used as the threshold for statistical significance. Significant fGSEA results were then collapsed into independent, or non-redundant gene sets using the collapsePathways function in the fGSEA package (24). The resulting output of significantly enriched gene sets were then ranked using the absolute normalized enrichment score (NES) for downstream analyses. All data visualizations were generated using either heatmap3 (27) or ggplot2 (28). All analyses were performed using R version 3.4.1 (29).

### 4. Cell deconvolution

Normalized microarray data was input into the xCell webtool (http://xCell.ucsf.edu/, accessed: 12/30/2021) to perform cell type enrichment analysis for the 64 immune and stroma cell types using the xCell signatures (N=64) (30). Results were downloaded and filtered using a significance threshold of p-value ≤ 0.2. The resultant significant cell type proportion data was visualized by boxplot using ggplot2 (28).

### 5. Animal wound model

Eight-week-old female Balb/c mice (Harlan, Inc., Indianapolis, IN) were anesthetized with intraperitoneal ketamine and xylazine injection. For skin wounds, the dorsal skin was shaved, and two one mm full thickness excisional wounds were made with a punch-biopsy instrument (Acu-Punch, Acuderm Inc., Ft. Lauderdale, FL). For oral wounds, one wound was made in the tongue using the same 1 mm punch biopsy instrument (1, 2, 4, 11, 31). At 6 hours, 12 hours, and Day 1 after injury, wounds and the surrounding tissue were removed with a 2 mm biopsy punch and stored in optimal cutting temperature (OCT) compound (Thermo Fisher Scientific, Waltham, MA). All animal procedures were approved by the University of Illinois Chicago Institutional Animal Care and Use Committee.

### 6. Quantification of mast Cells in mouse skin and oral wounds

To count the number of mast cells within the wound bed, 10 μm sections were prepared from frozen embedded wounds using a cryostat. Frozen sections were thawed and fixed for 1 hour in Carnoy’s fixative (60% ethanol, 30% chloroform, and 10% glacial acetic acid). Sections were stained for 2 hours at room temperature with 0.5% toluidine blue (Sigma, St. Louis, MO) in 0.5N HCl in PBS, followed by dehydration and xylene clearing. A cover glass was mounted on each sample with Cytoseal (Richard-Allan Scientific, Kalamazo, MI). Mounted sections were viewed under a 20x objective and mast cells were counted. Three sections per mouse were averaged. Wounds from five mice were counted. The total number of mast cells/mm^2^ of the wound sections was expressed in the result.

### 7. Data availability

Microarray gene expression data generated and analyzed in this study have been deposited at the NCBI Gene Expression Omnibus (GEO) database with the primary accession code GSE209609.

## Results

### 1. Human palate and skin wounds cluster both by tissue and time after injury

A wound healing microarray gene expression dataset of human palate and skin excisional wounds was generated from 18 subjects containing equal numbers of biological sex (9 female, 9 male), including five time points (0 hours, 6 hours, Day 1, Day 3, and Day 7), for a total of 96 samples (**Figure 1**). Initial visualization of the normalized data by principal component analysis (PCA) showed that the first two principal components account for 24.45% and 20.20% of the observed variance in the dataset, respectively, and these components distinguish the samples by time and tissue type (**Figure 2A**). Samples were then grouped by tissue and time point, a mean gene expression value was calculated for each gene within each assigned group, followed by computing a Pearson’s correlation of each group’s mean gene expression data, and hierarchical clustering was performed. Clustering showed that unwounded palate and skin groups were most similar to their respective Day 7 wound tissues (**Figure 2B**). Further, the acute response to injury groups (6 hours and Day 1) in both tissues clustered separately from the unwounded, Day 3, and Day 7 groups. Within the acute and late (Day 3 and Day 7) injury clusters, palate and skin were more similar to their respective tissue than time after injury. Taken together, palate and skin wounds separate first by an acute and late response to injury, and then by tissue, with overall transcriptomic profiles strongly correlating across all groups.

**Figure 1.**
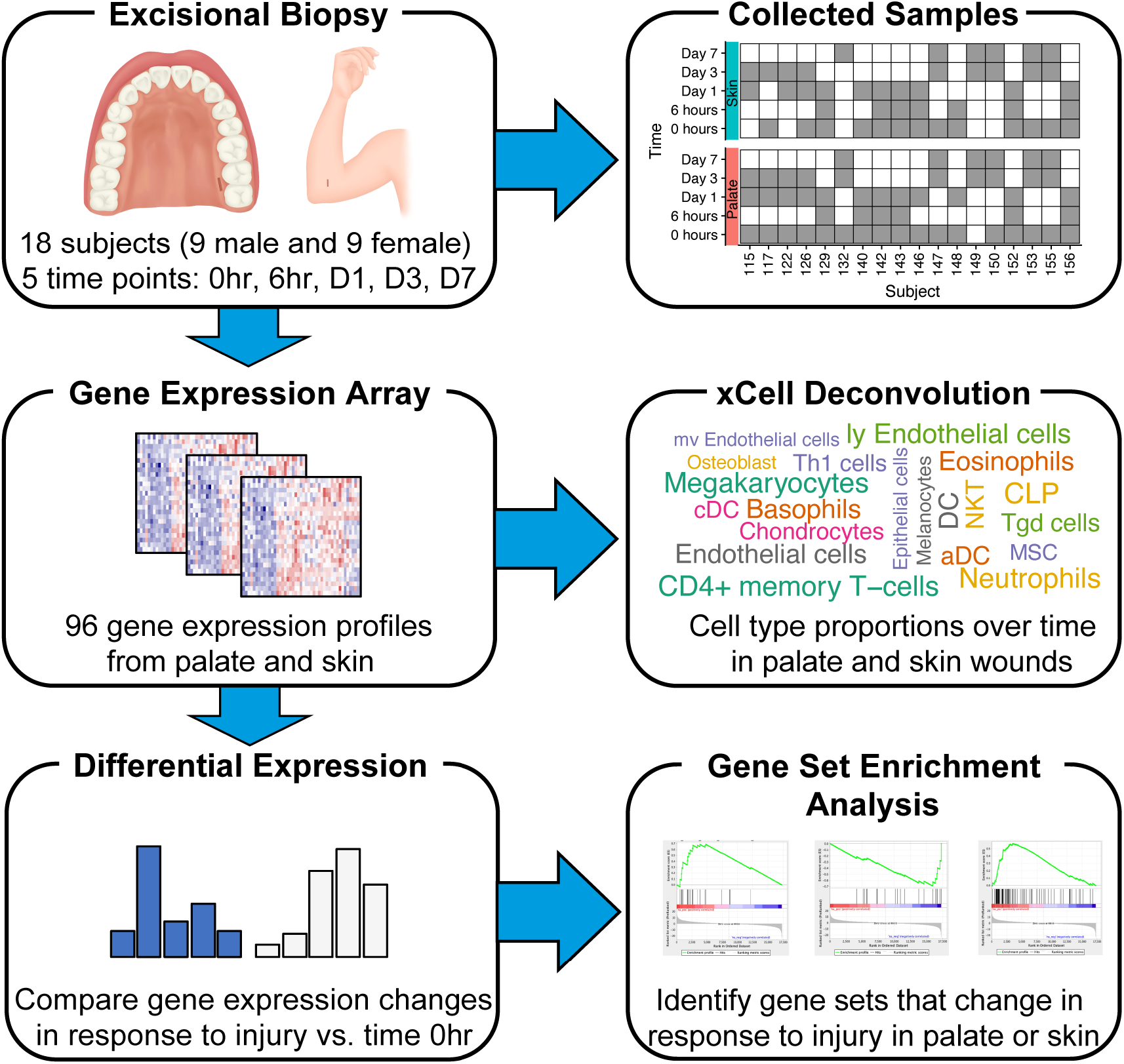
Study overview. Longitudinal excisional biopsies (1×5 mm) were placed in palate or skin and samples were collected (2×5 mm) at specific time points after injury. Each subject and the corresponding biopsies that were analyzed for this study are indicated by the grey boxes in the Collected Samples section. Microarrays were generated from RNA for a total of 96 samples. Differential expression analyses using LIMMA were done on a tissue-specific basis to compare each time point after injury to the respective unwounded tissue. Pre-ranked gene set enrichment analysis was done to identify functional changes in response to injury in palate and skin. A cell deconvolution analysis was performed to approximate cell type proportions in unwounded palate and skin and at each time point after injury.

**Figure 2.**
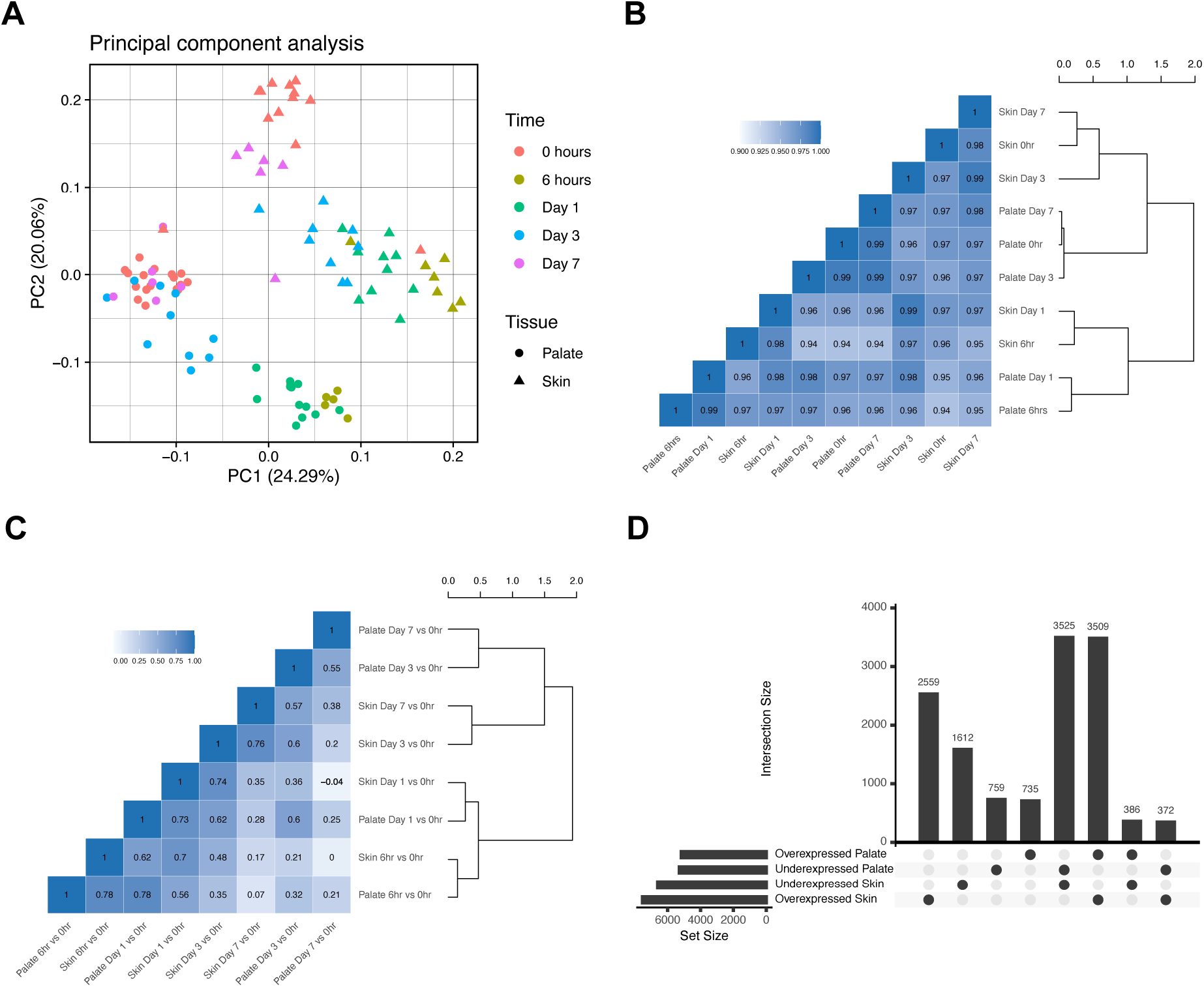
Comparative analysis of transcriptomic response to injury. A) Principal component analysis plot of microarray gene expression data. Each sample is represented by a point on the graph. Color represents time after injury with 0 representing unwounded tissue. The shape of each point represents tissue type. The x- and y-axis are the first and second principal components, respectively. B) Heatmap and dendrogram representing similarities of the mean gene expression profiles grouped by tissue and time point. A Pearson’s correlation coefficient based on the computed Euclidean distance matrix and hierarchical clustering using the complete agglomeration method were performed. Groups are denoted on both axes of the heatmap and adjacent to the dendrogram. Heatmap color and numbers within each square represent the correlation coefficients. C) Heatmap and dendrogram representing the similarities of t-statistics generated from the LIMMA differential expression contrasts. A Pearson’s correlation coefficient based on the computed Euclidean distance matrix and hierarchical clustering using the complete agglomeration method were performed. Groups are denoted on both axes of the heatmap and adjacent to the dendrogram. Heatmap color and numbers within each square represent the correlation coefficients. D) Upset plot displaying the overlap between gene expression changes over time in response to injury in palate and skin. The set size bar plot on the bottom left of the diagram represents the total number of unique genes significantly differentially overexpressed or underexpressed in palate or skin at any point in time after injury, with each row denoted by the group label. The intersection size and corresponding bar plot indicate the number of genes that are in each intersection. Solid black circles under each bar plot indicate which of the four groups are included in that specific intersection. Only intersections that demonstrated conserved or unique responses to injury in palate and skin were shown for clarity.

### 2. Transcriptomic response to injury in human skin is greater in magnitude than palate

We next sought to identify differentially expressed genes in response to injury in human palate and skin. To do this, a differential expression analysis was performed to compare the gene expression profiles of samples in each group at each point in time after wounding to their baseline unwounded tissue samples using the R package LIMMA. Briefly, LIMMA uses linear regression models to compare gene expression values for many genes across multiple samples. In this analysis, multiple two-group comparisons were performed by computing a moderated t-statistic using a Bayesian model to reduce the standard errors across the genes tested (21). The threshold for significance was an adjusted p-value FDR ≤ 0.05 based on the BH method (23). In palate, there were 7,118 (6 hours) and 7,779 (Day 1) statistically significant differentially expressed genes compared to unwounded palate (time 0) (**Table 1**). However, by Day 3 and Day 7, the number differentially expressed genes was dramatically reduced to 1,762 and 485, respectively. In the skin response to injury, there were 8,694 (6 hours), 8,384 (Day 1), 5,983 (Day 3), and 3,734 (Day 7) differentially expressed genes when compared to unwounded skin (time 0). The reduced number of differentially expressed genes compared to baseline in palate demonstrates that by as early as Day 3, palate wound transcriptomes are quite similar to unwounded palate transcriptomes. The large number of differentially expressed genes at each time point after injury in skin highlights the transcriptomic dissimilarity of skin wounds to their unwounded skin tissue at every time point examined.

**Table 1.**
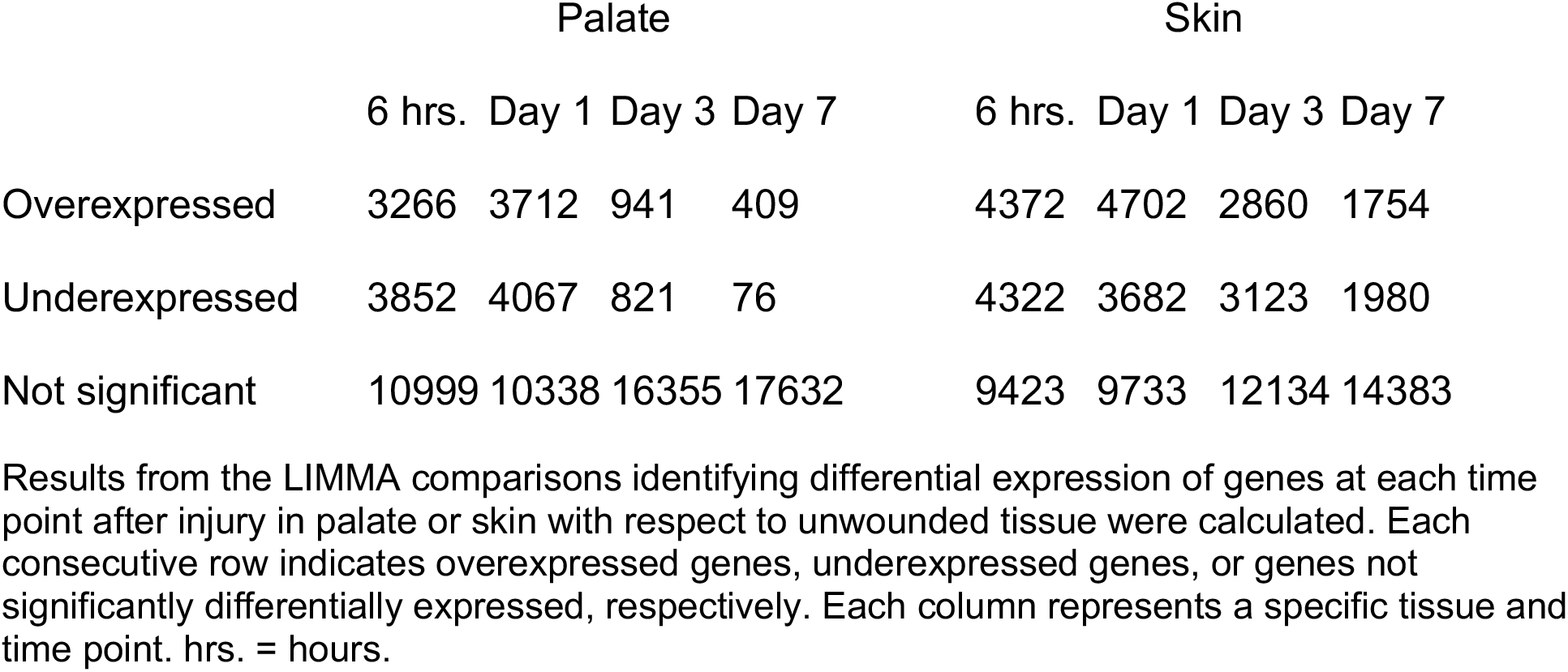
Differentially expressed genes over time in palate and skin.

|  | Palate |  |  |  | Skin |  |  |  |
| --- | --- | --- | --- | --- | --- | --- | --- | --- |
|  | 6 hrs. | Day 1 | Day 3 | Day 7 | 6 hrs. | Day 1 | Day 3 | Day 7 |
| Overexpressed | 3266 | 3712 | 941 | 409 | 4372 | 4702 | 2860 | 1754 |
| Underexpressed | 3852 | 4067 | 821 | 76 | 4322 | 3682 | 3123 | 1980 |
| Not significant | 10999 | 10338 | 16355 | 17632 | 9423 | 9733 | 12134 | 14383 |

Results from the LIMMA comparisons identifying differential expression of genes at each time point after injury in palate or skin with respect to unwounded tissue were calculated. Each consecutive row indicates overexpressed genes, underexpressed genes, or genes not significantly differentially expressed, respectively. Each column represents a specific tissue and time point. hrs. = hours.

To identify the level of conservation in the response to injury between palate and skin, a Pearson’s correlation and hierarchical clustering was performed using the t-statistics from each LIMMA contrast. Each contrast compares the response to injury at a specific time point to the respective unwounded tissue transcriptomic profile thereby controlling for any underlying baseline tissue-specific gene expression patterns. Interestingly, the acute response to injury time points (6 hours and Day 1) clustered together by time and not tissue type (**Figure 2C**). However, Day 3 and Day 7 samples clustered by tissue and not time point. The Pearson correlation values were reflected in the hierarchical clustering results. To compare the specific gene expression changes occurring over time in palate and skin, genes differentially expressed (FDR ≤ 0.05) at any time point after injury in each tissue were binned as overexpressed or underexpressed with respect to their unwounded tissue expression based on the t-statistic sign. In palate there were 5,171 genes overexpressed and 5,286 underexpressed after injury (**Figure 2D, Supplementary Data Files**). In skin there were 7,536 genes overexpressed and 6,606 underexpressed after injury. Of the total differentially expressed genes, 735 genes were overexpressed and 759 genes were underexpressed only in palate (palate-specific) in response to injury. In skin, 2,559 genes were overexpressed and 1,612 were underexpressed only in skin (skin-specific) in response to injury. There were 386 genes overexpressed in palate that were underexpressed in skin after injury, showing opposing regulation of these genes. Further, there were 372 genes overexpressed in skin and underexpressed in palate after injury. In regard to the conserved response, 3,509 genes were overexpressed after injury in both palate and skin, and 3,525 were underexpressed in both tissues. This analysis shows that while there is some degree of conservation in the transcriptomic changes in response to injury in palate and skin, a large number of genes are expressed in either a tissue-specific or opposing manner.

### 3. Temporal gene expression patterns in response to injury

Statistically significantly differentially expressed genes (FDR ≤ 0.05) were further filtered by absolute log2FC ≥ 1 for each comparison described above in palate and skin, which resulted in a total of 2,589 unique genes. The mean gene expression for each tissue and time point for these genes was then subjected to K-means clustering into five clusters. Genes within each cluster were plotted via heatmap together with a line plot to view the expression trends over time (**Figure 3**). Cluster 1 (369 genes) showed an increase in expression after injury in both palate and skin, with higher expression initially in the palate, but the skin later surpasses the palate expression, perhaps representing a later phase of the response to injury. Cluster 2 (218 genes) had increased expression in palate when compared skin at all time points, with clusters from both tissues following the same trend. There is an initial decrease in expression at 6 hours and Day 1, slowly returning to and then exceeding baseline expression by Day 7. Cluster 3 (573 genes) is most representative of the acute response to injury, as increases in gene expression peak at 6 hours and slowly return to baseline over time. Cluster 4 (966 genes) genes were underexpressed at 6 hours after injury, and slowly increased to their original expression over by Day 7. Cluster 5 (463 genes) genes have increased expression at 6 hours and Day 1 in both palate and skin, with a peak expression in palate and skin at 6 hours. These clustering results demonstrate that a majority of the most highly differentially expressed genes tend to follow similar trends in both tissues over the first seven days of wound healing.

**Figure 3.**
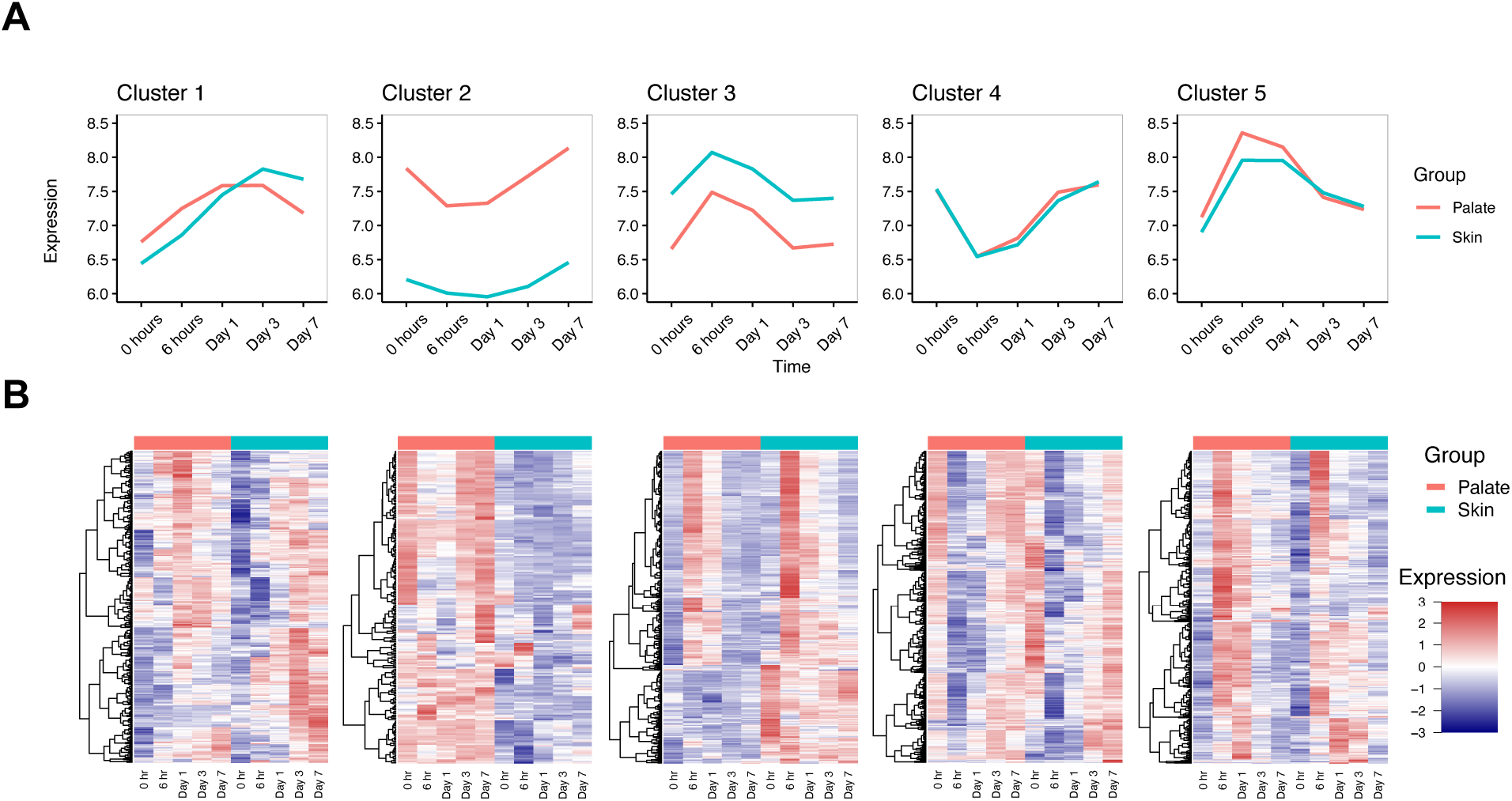
Transcriptomic trends in tissue repair in human palate and skin. A) Differential expression results for each LIMMA comparison were filtered for significance by an adjusted p-value ≤ 0.05 and an absolute log2FC ≥ 1. The average gene expression for each group (based on tissue and time point) were used to perform K-means clustering (k=5). For each cluster, a line plot of the mean expression of genes (y-axis) at each time point representing hours after injury (x-axis) was made. Unwounded tissue is represented by time 0. Clusters are denoted by titles above each plot, with orange lines representing palate and green lines representing skin. B) Heatmaps of each of the five identified clusters using the mean gene expression for each group (based on tissue and time point) were generated. Rows are hierarchically clustered using Euclidean distance and the Complete agglomeration method. Unwounded tissue is represented by time 0. Each row represents a gene, and each column represents the average expression of that gene for the specified group. Tissue type is represented on the top of the columns by palate in orange and skin in green.

### 4. Gene set enrichment analysis of palate and skin response to injury

Gene Set Enrichment Analysis (GSEA) (32) was performed using pre-ranked t-statistics from each LIMMA differential expression comparison to identify biological processes that change in response to injury in human palate and skin at each time point. An adjusted p-value (FDR ≤ 0.05) was used as the threshold for statistical significance. Of the 1,004 gene sets that were enriched in at least one fGSEA comparison, there were 348 unique gene sets enriched only in the palate response to injury compared to baseline tissue expression, 321 unique gene sets enriched only in the skin response to injury, and 335 unique gene sets enriched in both palate and skin. The top 25 overlapping gene sets enriched in at least one time point after injury in both palate and skin were ranked by absolute NES and plotted (**Figure 4**). The top five gene sets were: 1) Hallmark epithelial mesenchymal transition, 2) GO ribosome biogenesis, 3) Hallmark tumor necrosis factor alpha (TNFα) signaling via nuclear factor kappa-light-chain-enhancer of activated B cells (NFκB), 4) GO extracellular structure organization, and 5) GO ribonucleoprotein complex biogenesis. Next, gene sets enriched only in skin or palate at multiple time points were investigated. In the palate-specific gene sets, there were 51, 12, and 10 gene sets enriched in two, three, or four time points after injury, respectively (**Table 2**). Of the skin-specific gene sets, there were 58, 15, and 1 gene sets enriched in two, three, or four time points after injury, respectively. Palate-specific and skin-specific gene sets were ranked by absolute NES and the top 25 were plotted (**Figure 5**). The top three palate-specific gene sets were: 1) GO endothelium development, 2) GO positive regulation of chemotaxis, and 3) GO cellular response to vascular endothelial growth factor (VEGF) stimulus. The top three skin-specific gene sets were: 1) KEGG extracellular matrix interaction, 2) GO response to Type I interferon, and 3) GO translational termination.

**Figure 4.**
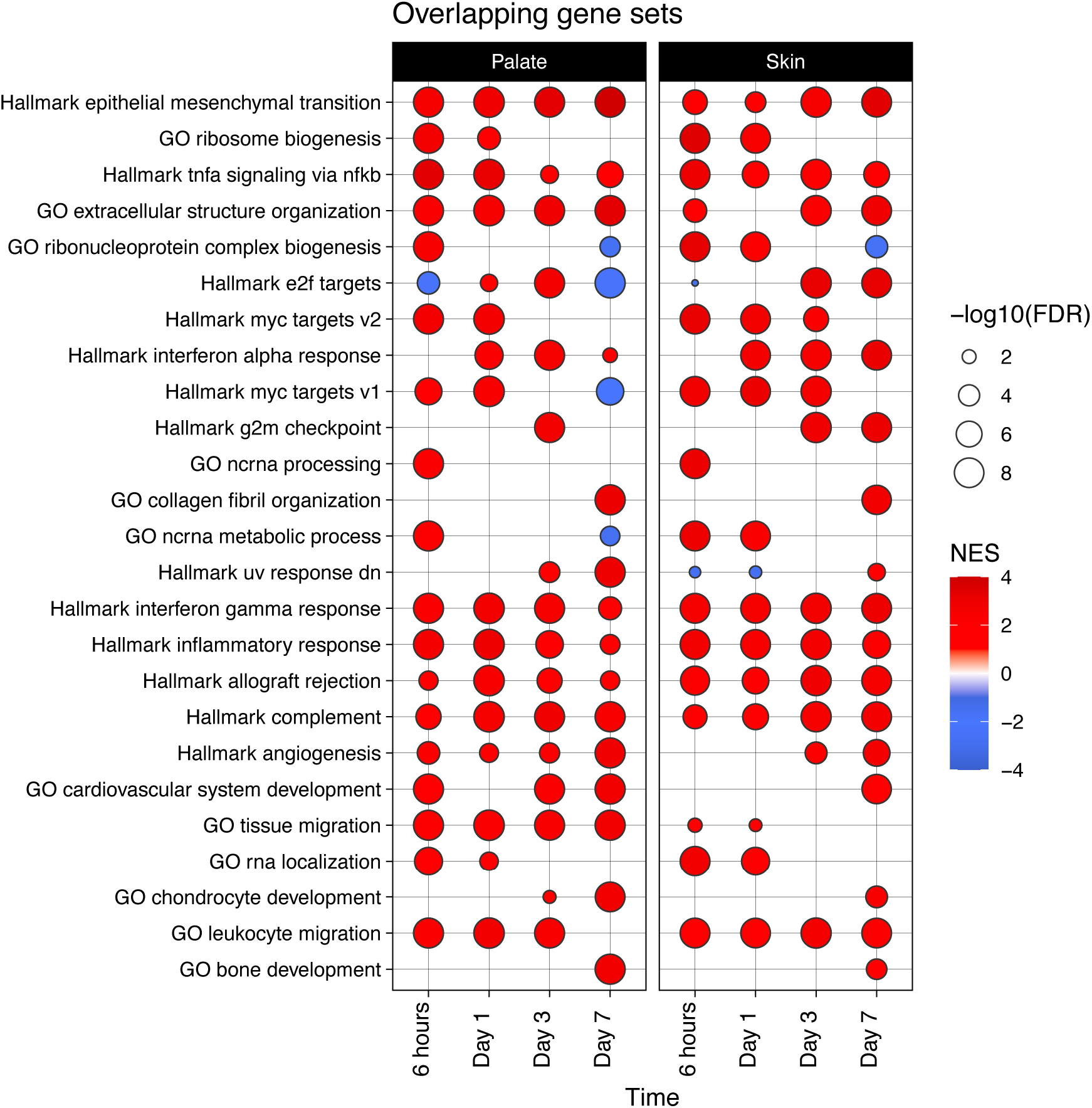
Overlapping gene sets enriched in both palate and skin. Bubble plot representing the gene sets enriched in both palate and skin at a specific time point after injury when compared to the respective uninjured tissue. The left panel represents palate, and the right panel represents skin, with each column representing the time point after injury. Rows are labeled by the enriched gene set name and are ranked in descending order of their NES. Presence of a circle indicates the gene set was enriched at that time point in that tissue. Color indicates the NES and bubble size indicates the statistical significance -log10(FDR).

**Figure 5.**
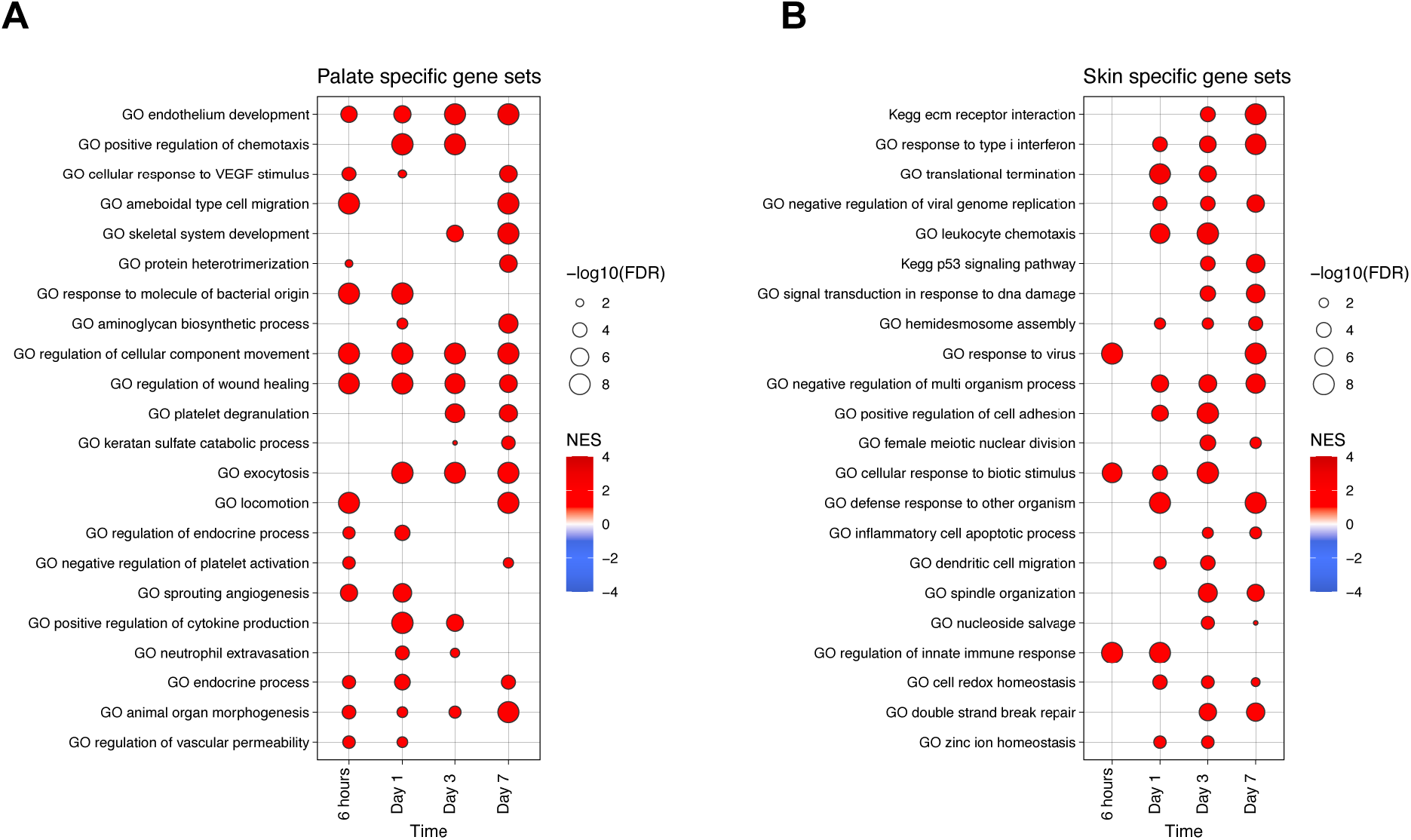
Tissue-specific gene sets enriched at multiple time points after injury. Bubble plot representing the gene sets enriched specifically in A) palate or B) skin at multiple time points after injury when compared to the respective uninjured tissue. The left panel represents palate, and the right panel represents skin, with each column representing the time point after injury. Rows are labeled by the enriched gene set name and are ranked in descending order of their NES. Presence of a circle indicates the gene set was enriched at that time point in that tissue. Color indicates the NES and bubble size indicates the statistical significance -log10(FDR).

**Table 2.** Frequency table of enriched gene sets across comparisons.

| Frequency | 1 | 2 | 3 | 4 | 5 | 6 | 7 | 8 |
| --- | --- | --- | --- | --- | --- | --- | --- | --- |
| Palate specific | 275 | 51 | 12 | 10 | 0 | 0 | 0 | 0 |
| Skin specific | 247 | 58 | 15 | 1 | 0 | 0 | 0 | 0 |
| Overlap | — | 89 | 91 | 51 | 36 | 33 | 23 | 12 |

Statistically significantly enriched gene sets for each LIMMA comparison from the fGSEA analysis were counted. As there were 8 comparisons, one for each tissue and each time point after injury (4 per tissue), a gene set could be enriched up to a maximum of 8 times. Each row indicates the unique gene sets that were enriched only in palate (Palate specific), only in skin (Skin specific) or gene sets enriched in both tissues (Overlap). Each column represents the number of times, or frequency, a particular gene set was enriched.

Gene sets that were uniquely enriched in a palate- or skin-specific manner and enriched at only one time point after injury were also investigated. The top 25 gene sets for each time point were ranked by absolute NES and plotted. In palate at 6 hours, gene sets related to catabolism of branched chain amino acids, propanoate metabolism and processes related to DNA repair and regulation of centriole replication were negatively enriched (**Figure 6**). Positively enriched gene sets at 6 hours in palate included GO positive regulation of epithelial cell migration, GO response to epidermal growth factor, GO regulation of anoikis, GO regulation of extrinsic apoptotic signaling, and GO endoplasmic reticulum unfolded protein response. Gene sets enriched at Day 1 included regulation of platelet activation and aggregation, granulocyte migration, establishment of endothelial barrier, interleukin (IL) signaling and production, and the extracellular-signal-regulated kinase (ERK) ERK1 and ERK2 cascade. Enriched gene sets at Day 3 were related to many immune processes such as leukocyte chemotaxis and migration, antigen processing and presentation, GO wound healing, GO negative regulation of immune response, and GO negative regulation of humoral immune response. By Day 7, many of the top 25 upregulated gene sets in palate were related to developmental or morphogenic processes including GO tube morphogenesis, GO connective tissue development, and GO mesenchyme morphogenesis.

**Figure 6.**
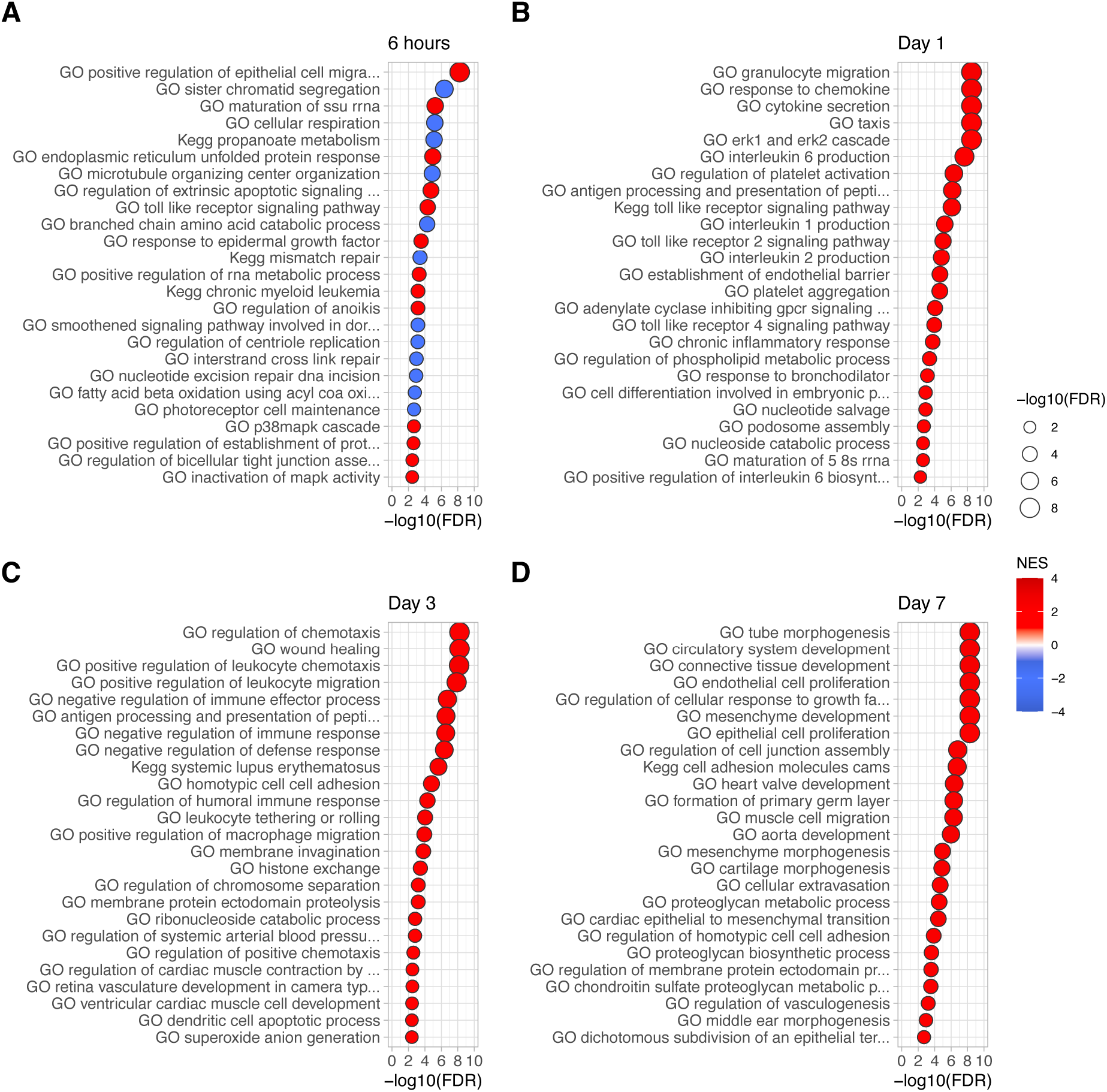
Time-point specific enriched gene sets in palate. Bubble plots representing the top 25 gene sets enriched uniquely in palate and only at the indicated time point after injury when compared to the respective uninjured tissue. Rows are labeled by the enriched gene set name and are ranked in descending order of their adjusted p-value. Bubble color indicates the NES, x-axis and bubble size represent the -log10(FDR). Each plot represents a time point after injury either A) 6 hours, B) Day 1, C) Day 3, or D) Day 7.

In contrast to palate, the top 25 enriched gene sets in skin at 6 hours were involved in DNA transcription and elongation, mRNA processing, mitochondrial protein import and RNA processing, Interleukin-1 (IL-1) mediated signaling pathway, and phosphorylated ERK mediated unfolded protein response (**Figure 7**). At Day 1 in skin, enriched gene sets included KEGG proteasome, cellular processes related to the nuclear export and nuclear pore organization, protein export, metabolic processes, and GO regulation of neutrophil migration. Day 3 enriched gene sets included GO cell killing, GO innate immune response, GO Interleukin-12 (IL-12) production, and GO glycosyl compound catabolic process. By Day 7 the enriched gene sets in skin involved T-cell migration and chemotaxis, Interferon alpha (IFNα) production, and other immune cell processes. Taken together, there are many conserved processes that occur in both palate and skin in response to injury that are largely related to a generalized immune response and ECM reorganization. However, the palate response appears to have an abrupt immune response that ends to allow for regenerative processes as early as Day 7. In the skin there appears to be an immune response that is still present at Day 3 and Day 7.

**Figure 7.**
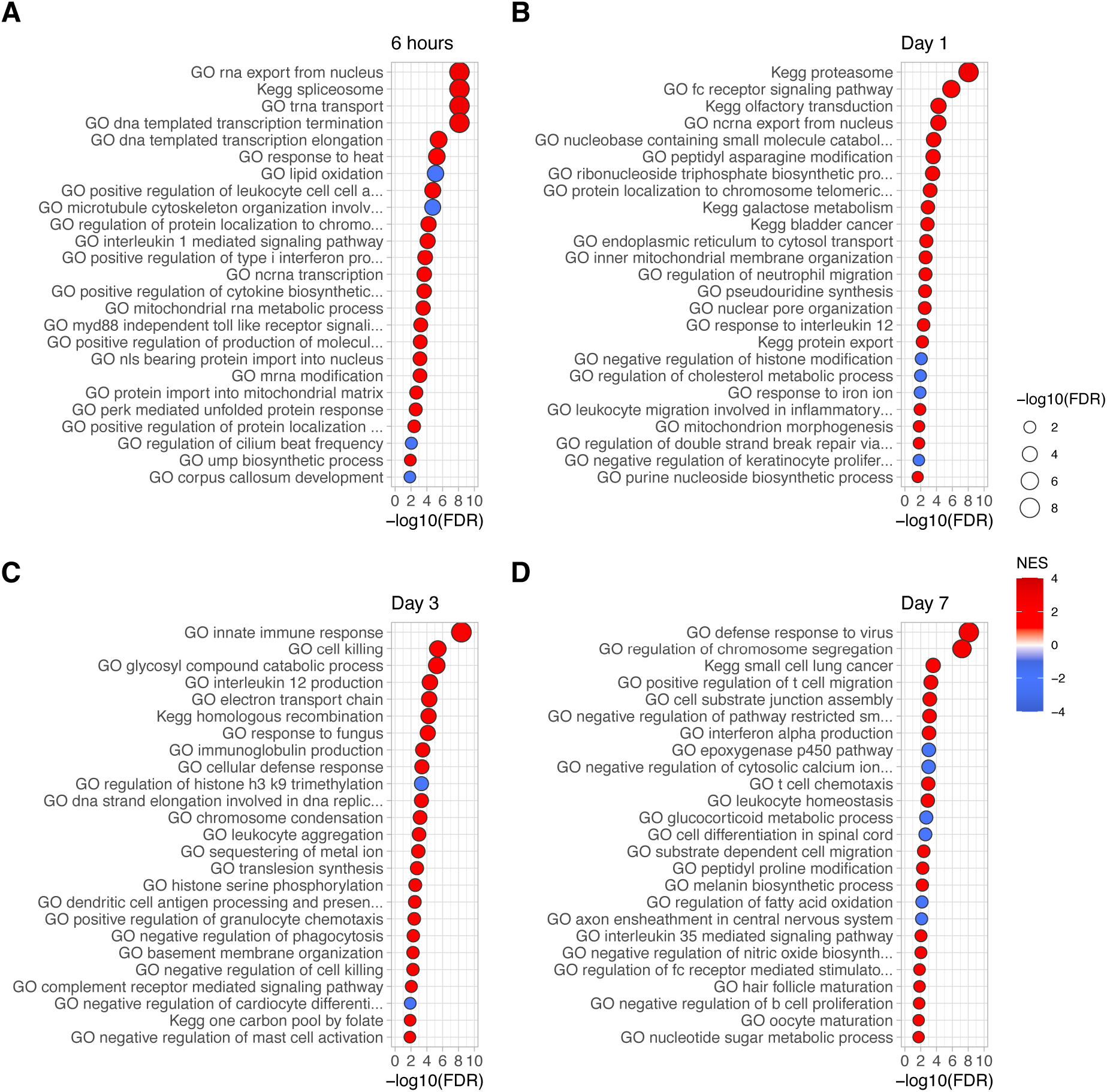
Time-point specific enriched gene sets in skin. Bubble plots representing the top 25 gene sets enriched uniquely in skin and only at the indicated time point after injury when compared to the respective uninjured tissue. Rows are labeled by the enriched gene set name and are ranked in descending order of their adjusted p-value. Bubble color indicates the NES, x-axis and bubble size represent the - log10(FDR). Each plot represents a time point after injury either A) 6 hours, B) Day 1, C) Day 3, or D) Day 7.

### 5. Cell type distribution changes over time in palate and skin wounds

Previous studies suggest that cell type distribution varies between palate and skin healing over time and could be responsible in part for the observed differential healing outcomes in palate and skin (1). To investigate this, a cell deconvolution analysis was performed using the microarray gene expression data and the xCell algorithm to approximate the proportion of cell types in palate and skin at each of the five experimental time points. The xCell signature dataset was used (N=64 cell types) as it includes many different stromal and immune cell types and was most similar to the expected cellular makeup in these tissues (30). This resulted in the identification of a higher proportion of dendritic cells (DC) in skin at each time point after injury when compared to palate (**Figure 8**). There was also an observed increased proportion of mast cells in skin after injury, with more overall in unwounded skin than palate. This correlates well with a histologic comparison of mast cell numbers in oral and skin murine wounds over the first 24 hours *in vivo* (**Supplementary Figure 1**). Neutrophils rose significantly in the first 6 and 24 hours after injury in both tissues, then declined significantly by Day 3 in palate while remaining high in skin. There were also more natural killer T-cells (NKT) and mesenchymal stem cells (MSC) after injury in skin than palate. However, endothelial cells and keratinocytes showed higher proportions after injury in palate than skin. The complete set of all 64 cell type proportions were included in the initial analysis (**Supplementary Figure 2**). This data demonstrates that there are potentially very significant differences in wound cell type populations over the seven days post injury in human palate and skin wounds.

**Figure 8.**
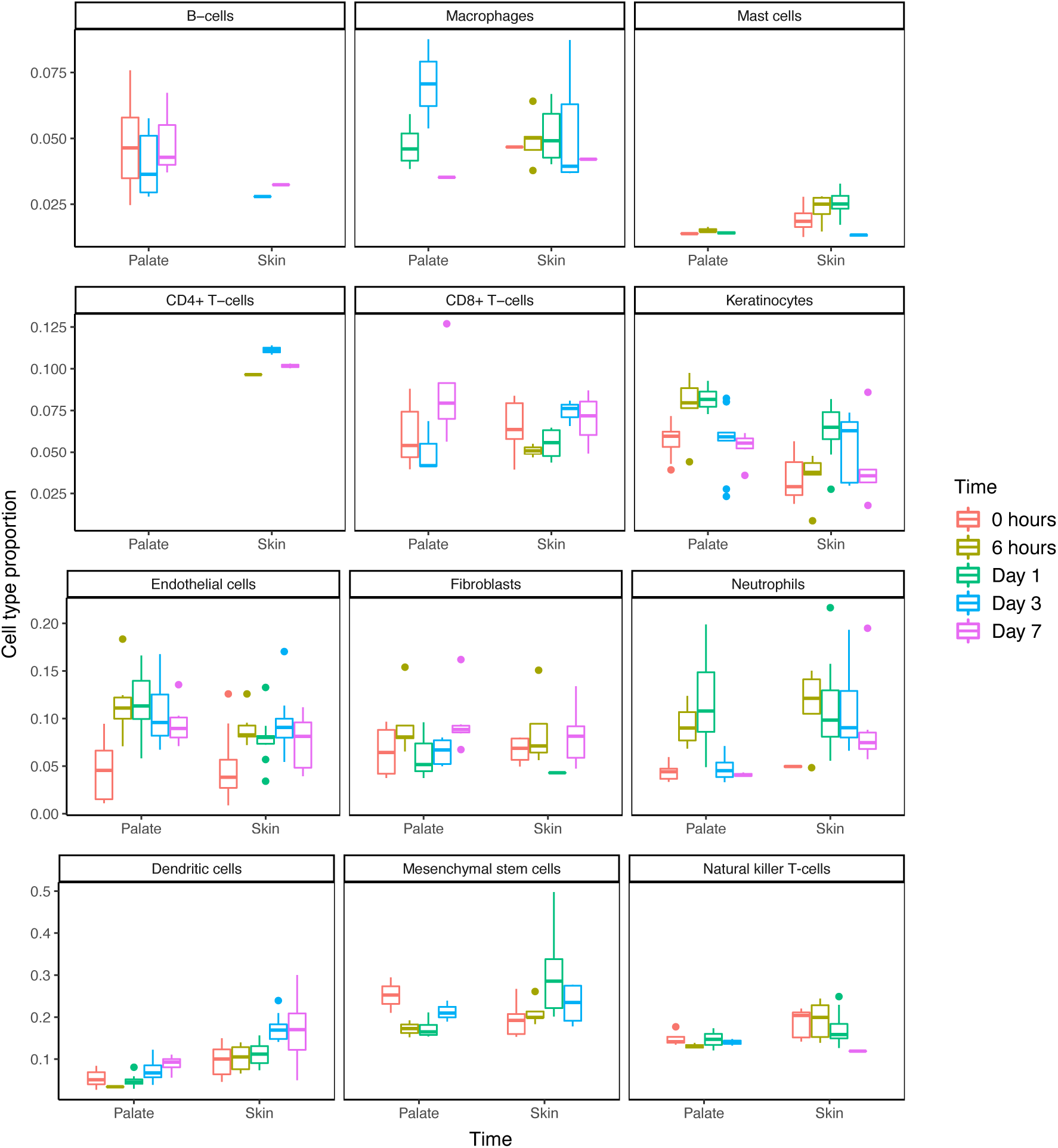
Cell deconvolution analysis of cell type proportions in tissue injury. Results from the xCell deconvolution analysis were first filtered for statistical significance (p-value ≤ 0.2). Boxplots representing the cell type proportion (y-axis) at each point in time after injury in palate or skin (x-axis) were generated for each cell type (denoted by panel title). Colors represent the time point after injury, with 0 representing unwounded tissue. Outlier samples are represented by circles on the plots. Boxplots that do not appear for a specific tissue and time point indicate that the results from the xCell analysis did not reach statistical significance.

## Discussion

Tissue repair in response to injury can lead to a relatively minimal loss in functional tissue, or conversely, a fibrous collagen laden scar. Most adult tissues in humans have limited regenerative capabilities and are prone to a scarring outcome when injured. The oral cavity responds to tissue injury with minimal scarring where the precise mechanisms responsible for this outcome have yet to be fully elucidated. To better understand the differences specifically in human tissue repair and regeneration, we generated a wound healing microarray gene expression dataset of human palate and skin excisional biopsies, analyzed the transcriptomes of 96 samples from 18 patients across five time points, and performed an in-depth computational analysis including differential expression and gene set enrichment analyses to characterize the first seven days after injury. These data represent a unique and significant resource for the wound healing field.

One open question in the field is whether the skin response to injury is simply a delayed version of the oral response to injury. To investigate this, we looked at wound samples over time in each tissue type and compared the similarity of transcriptomic profiles. This showed that the acute transcriptional response to injury (6 hours and Day 1) in palate and skin cluster separately from the respective unwounded palate and skin tissues. To ensure that tissue-specific baseline expression was not swamping out the response to injury signal, the t-statistics of the LIMMA comparisons of each time point after injury versus unwounded tissue were grouped and correlated. This analysis demonstrated that the acute transcriptomic response to injury appears to be more similar between palate and skin, while the late time points become more tissue specific. Taken together, these analyses demonstrate that, at least within the first 7 days of wound healing, the skin is not simply a delayed version of the palate response to injury.

A more quantitative examination of the transcriptomic differences in response to injury in palate and skin was performed by analyzing the differential expression data. These data showed that there is a transcriptomic response to injury in skin that is greater in magnitude than the palate response to injury. A majority of the differentially expressed genes (7,034) were overexpressed or underexpressed in the same manner in both palate and skin, while smaller subsets of genes differentially expressed after injury were unique to palate (1,494), to skin (4,171), or had opposing expression patterns (758) in palate and skin injuries. The magnitude of differentially expressed genes in human skin over time was ∼2.79 fold that seen in the palate injury. This is similar to what was observed in our previous studies comparing mouse tongue and skin wounds, where skin had more significantly differentially expressed genes over the 10 day response to injury than tongue (4). Overall, this quantitative analysis highlights both the similarities and differences in the transcriptomic response to injury in human palate and skin.

The fGSEA was able to attribute biologically meaningful information to the differential expression data. From the quantitative analysis of differentially expressed genes, it was clear that there were both conserved, or generalized wound responses, and tissue-specific responses to injury, as has been observed in tissue repair studies in other organisms (33). The fGSEA results showed that this generalized wound response involves gene sets and processes including the epithelial mesenchymal transition, migration of multiple immune cell types, extracellular matrix organization, and many hallmark inflammatory gene sets enriched at multiple time points after injury in both palate and skin (**Figure 4**). Gene sets enriched in multiple time points after injury in only palate or skin highlight the tissue-specific responses to injury. In palate, GO endothelium development, GO cellular response to VEGF stimulus, GO sprouting angiogenesis, and GO regulation of vascular permeability were all upregulated at multiple time points. Previous studies have highlighted that vessel density changes are greater in skin after injury when compared to the respective unwounded tissue vessel densities (2). However it is also important to note that at baseline, the percent vascularization as marked by cluster of differentiation 31 (CD-31) to wound area (CD-31 positive area / Total area measured) was ∼2.5% in mucosa versus ∼0.5% in skin, indicating the high degree of vascularization in oral mucosa at baseline (2). The top gene sets overlapping at multiple time points that were skin-specific were more closely related to the immune response. These included GO dendritic cell migration, GO leukocyte chemotaxis, GO inflammatory cell apoptotic process, and GO regulation of innate immune response, which demonstrates the sustained inflammatory response in skin. The GO hemidesmosome assembly gene set was also enriched at every time point in skin injury except 6 hours, highlighting the importance for skin to restore barrier function.

The enriched gene sets unique to one time point and tissue highlight the dynamic changes that occur over brief periods of time in a tissue-specific manner. In the acute 6-hour response to injury in palate enriched gene sets were related to re-epithelialization, preventing cell-detachment induced cell death, regulation of extrinsic apoptotic signaling, the unfolded protein response, and response to epidermal growth factor (**Figure 6**). The top gene sets at Day 1 in palate included the most GO gene sets related to the immune response, including toll-like receptor signaling gene sets, IL-1 and Interleukin-2 (IL-2) production, and regulation of Interleukin-6 (IL-6) biosynthesis. There was also enrichment of ERK1 and ERK2 cascade, which has many downstream effects on cell survival, proliferation, and migration in wound healing (14). The acute response of preventing cell death, in concert with cell stress responses, may promote the regenerative response observed in palate at later time points. Indeed, by Day 7 the top gene sets overexpressed in palate were GO morphogenesis and development processes. This is in contrast to skin, where the acute response to injury in the first 6 hours and Day 1 included many enriched GO gene sets related to DNA transcription, metabolic gene sets, and immune gene sets (**Figure 7**). By Day 3 the skin-specific enriched gene sets were largely inflammatory, which is likely due to the large inflammatory cell infiltrate seen in skin wounds at this time. Unlike the developmental processes seen in palate at Day 7, the only top ranked skin-specific gene set related to development was GO hair follicle maturation.

An acute tissue response to injury that prevents cell detachment induced cell death or other forms of apoptotic cell death may be a key player in setting the stage for a more long-term regenerative response of the palate. This, combined with very specific and short duration cytokine and immune responses could potentially curtail what would otherwise be a significant inflammatory response such as the one observed in skin. It is clear that both skin and palate tissues have strong immune signaling responses that involve cell chemotaxis and migration, IFNα and IFNγ responses, and TNFα signaling via NFκB. What remains unclear is how the tissue-specific enriched gene sets related to immune cell signaling processes are involved in the overall response to injury, and whether this is responsible for the differential healing outcomes in palate and skin. The cell deconvolution data identified specific populations of immune cells that were in higher proportion in skin than palate, including CD4+ T-cells, dendritic cells, mast cells, and natural killer T-cells. Mast cells have been shown to be required for normal healing in mouse wounds, but also contribute to scar formation in fetal wound models (34-36). Other studies have shown that blockade of mast cell activation can reduce cutaneous scar formation (37). The cell deconvolution analysis was primarily an exploratory approach to gain new insights. While differential immune responses have been previously reported between oral and skin wounds, the cell type proportions inferred using the xCell enrichment analysis will need further validation through cell sorting and single cell sequencing experiments (1).

In summary, we observed that during repair, the acute (6 hour and Day 1) response to injury is most similar between palate and skin, and that underlying tissue-specific expression profiles should be corrected for when performing between-tissue comparisons of the wound response. Skin was shown to have a greater magnitude of transcriptomic changes when compared to palate injury over time. We also identified biological processes shared between palate and skin across multiple time points after injury, highlighting the idea that certain aspects of the wound response to injury are conserved across tissues in an organism (33). However, later time points in palate showed a more regenerative phenotype when compared to the immune activated skin, again highlighting that there may be a conserved acute wound response and a second tissue-specific regenerative response (33).

One important limitation of the current study is the use of whole tissue samples for generation of the microarray data. Future investigations of oral and skin wounds using single cell technologies and spatial transcriptomics have the potential to link these observed phenotypes to specific cellular populations. Further investigation on what is driving these transcriptomic changes has the potential to identify core regulators of the various complex processes that underly the wound healing response. It has previously been proposed that taking a network approach could inform our current understanding of the molecular mechanisms of wound healing and could lead to new ways to control or optimize the process for better healing outcomes (38).

## Author Contributions

TRL, LAD, PTM, KG, and LC conceived and designed the experiments. PTM collected the human samples; TRL, MS, LC, and ZJF performed the experimental procedures. TRL performed the data analysis, prepared the figures, and drafted the manuscript, which was edited by LAD, KG, and LC. All authors reviewed the data and approved the manuscript.

## List of abbreviations

BH: Benjamini & Hochberg
CD-31: Cluster of differentiation 31
CDF: Chip description file
DC: Dendritic cells
EN1: Engrailed-1
ERK: Extracellular-signal-regulated kinase
FDR: False discovery rate
fGSEA: fast Gene Set Enrichment Analysis
GEO: Gene Expression Omnibus
GO: Gene Ontology
GSEA: Gene Set Enrichment Analysis
IFNα: Interferon alpha
IL: Interleukin
IL-1: Interleukin 1
IL-2: Interleukin 2
IL-6: Interleukin 6
IL-12: Interleukin 12
IRB: Institutional Review Board
KEGG: Kyoto encyclopedia of genomes and genes
LIMMA: Linear models for microarray data
log2FC: log2 fold change
Mm: millimeters
MSC: Mesenchymal Stem Cells
MSigDB: Molecular Signatures Database
NES: Normalized Enrichment Score
NFκB: nuclear factor kappa-light-chain-enhancer of activated B cells
NKT: Natural Killer T-cells
OCT: Optimal cutting temperature
PCA: Principal Component Analysis
PITX1: Paired like homeodomain 1
RMA: Robust multichip averaging
SOX2: SRY-box transcription factor 2
TNFα: Tumor necrosis factor alpha
UIC: University of Illinois Chicago
VEGF: Vascular endothelial growth factor

