## Supplemental Figures 1 and 2 for "Transcriptional changes in human palate and skin healing"

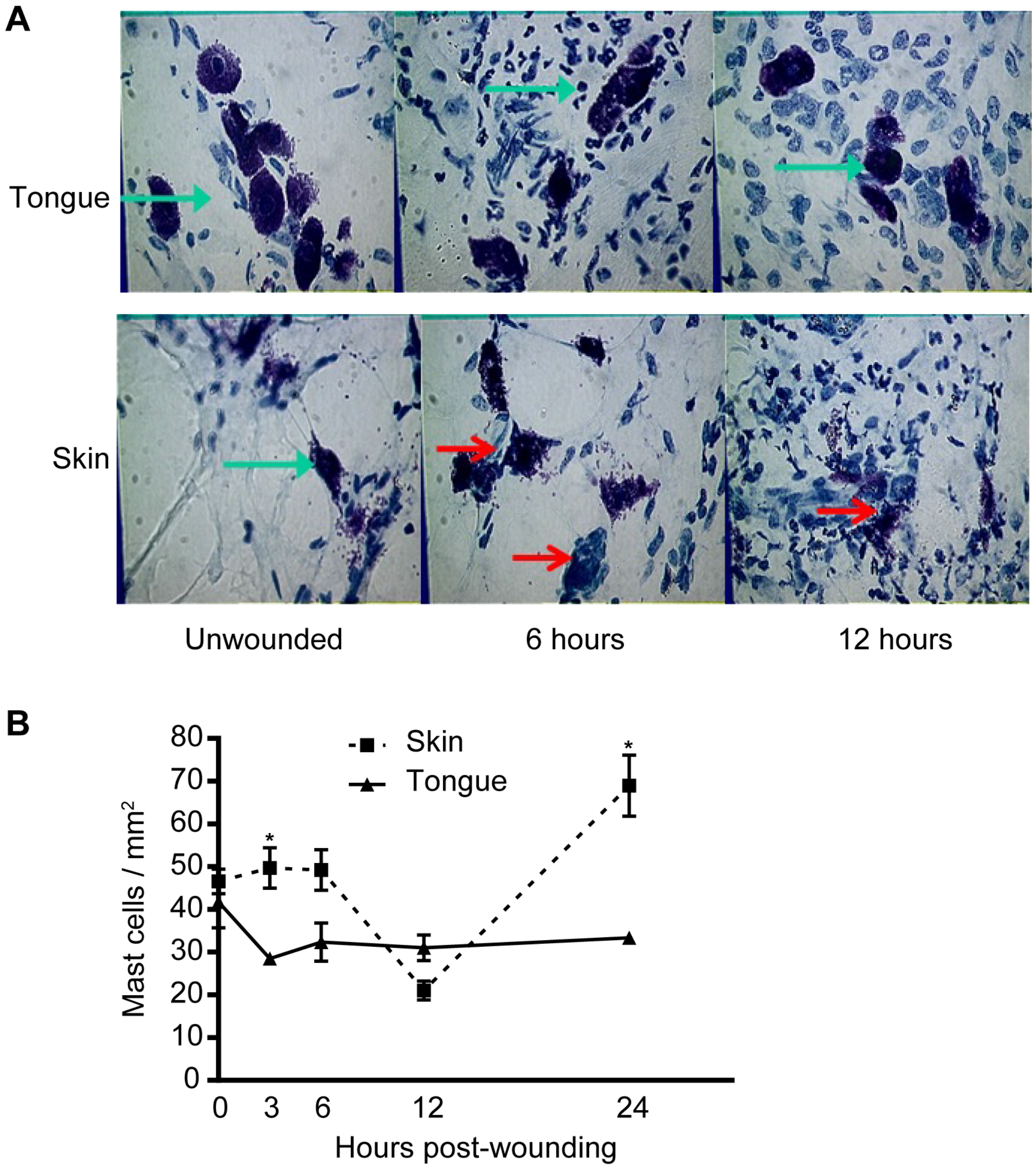


**Supplemental Figure 1. Histological analysis of mast cells in oral and skin wounds**.
A) Mast cells were visualized by toluidine blue staining of frozen sections from 1 mm tongue wounds (top panels) or 1 mm skin wounds (bottom panels). This method stains mast cell granules. Granulated mast cells stain dark purple (green arrows). Red arrows delineate partially degranulated mast cells. Unwounded tissue from both sites contained granulated mast cells. Representative sections from 6- and 12-hours post-wounding demonstrate that oral wounds contain many granulated mast cells (green arrows) while skin wounds contain partially degranulated mast cells (red arrows). B) Comparison of mast cell numbers in skin and oral wounds. Total mast cells numbers were counted using toluidine blue staining of tissue sections from skin wounds (squares) or oral

wounds (triangles). The total number of mast cells per square millimeter was counted. Three sections per mouse were averaged. Data is represented as total mast cells per area of the wound (n=5). *$p \leq 0.05$ by 2-way ANOVA followed by a Bonferroni post-test.

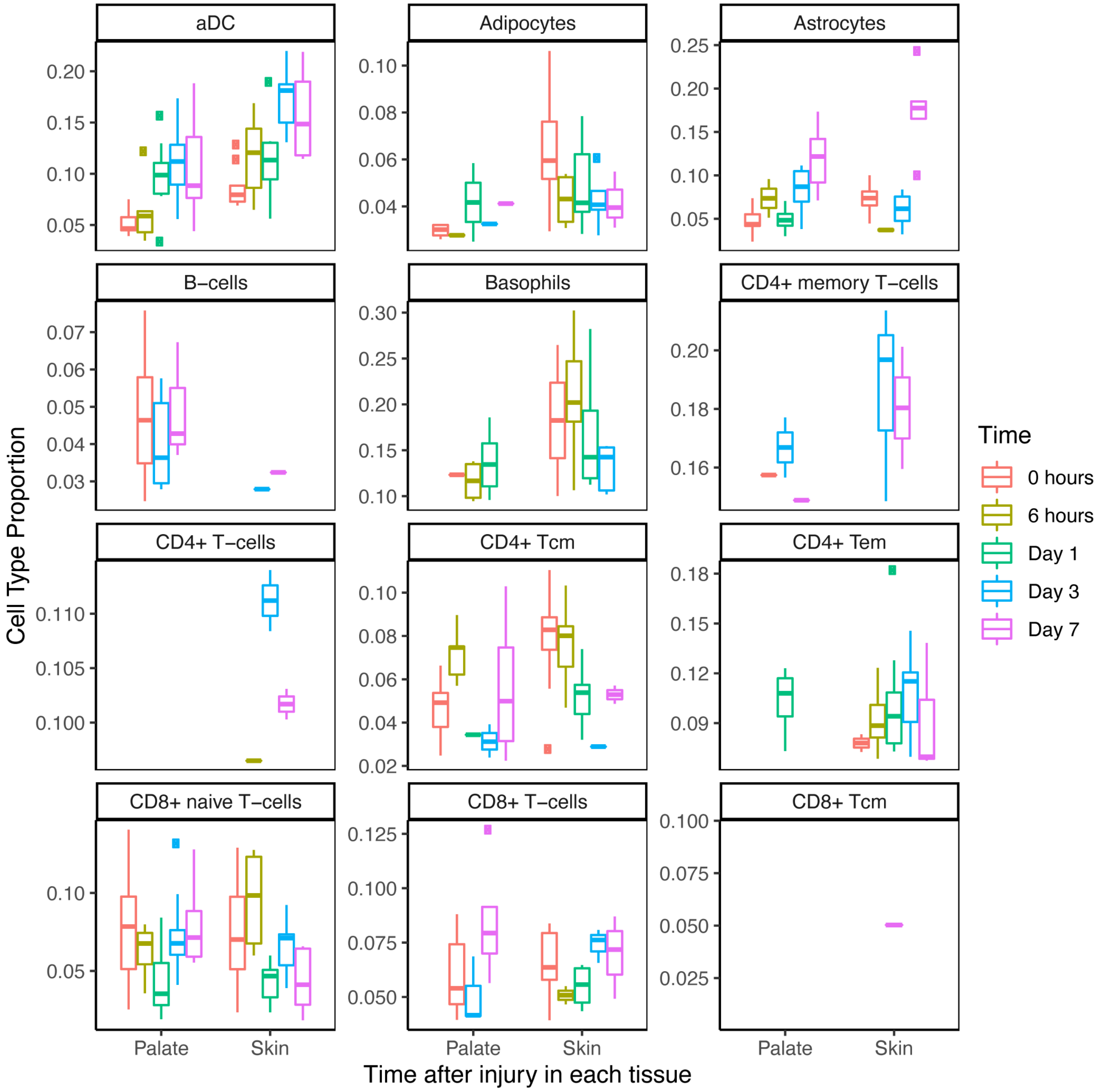

aDC
Adipocytes
Astrocytes
B−cells
Basophils
CD4+ memory T−cells
CD4+ T−cells
CD4+ Tcm
CD4+ Tem
CD8+ naive T−cells
CD8+ T−cells
CD8+ Tcm
Cell Type Proportion
Palate
Skin
Time after injury in each tissue
Time
0 hours
6 hours
Day 1
Day 3
Day 7

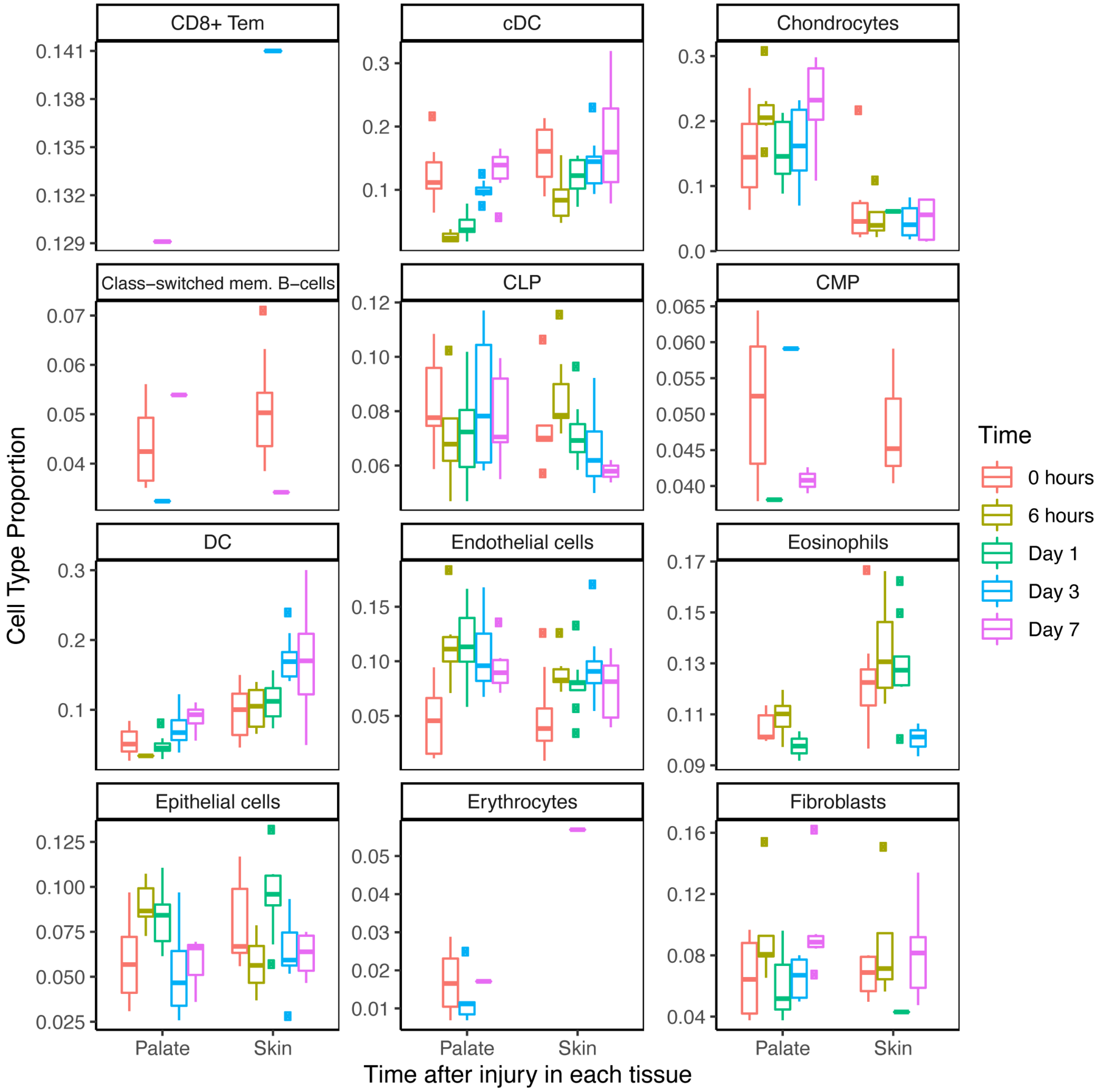
CD8+ Tem
cDC
Chondrocytes
Class−switched mem. B−cells
CLP
CMP
DC
Endothelial cells
Eosinophils
Epithelial cells
Erythrocytes
Fibroblasts
Cell Type Proportion
Time after injury in each tissue
Palate
Skin
Time
0 hours
6 hours
Day 1
Day 3
Day 7

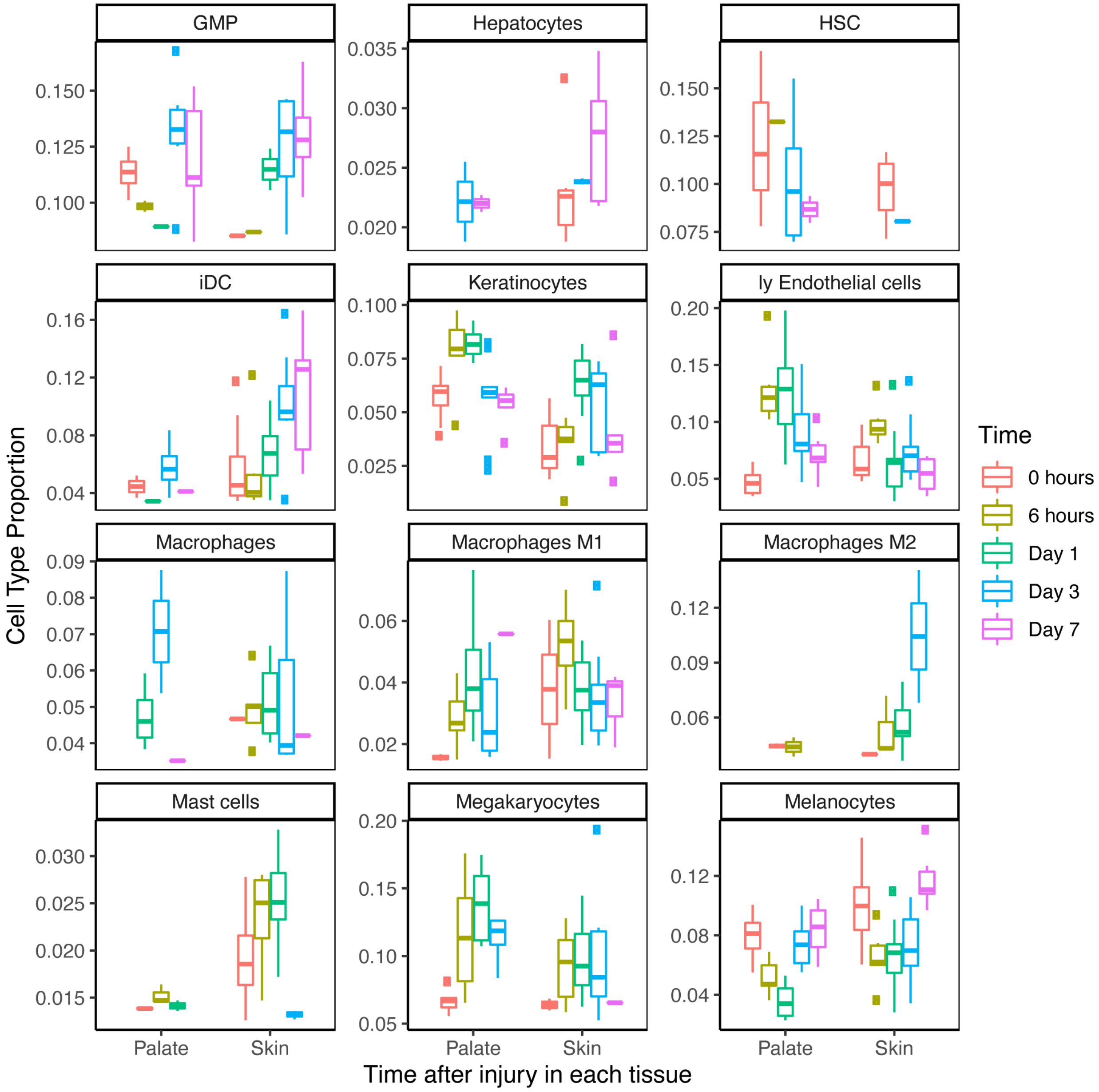
GMP
Hepatocytes
HSC
iDC
Keratinocytes
ly Endothelial cells
Macrophages
Macrophages M1
Macrophages M2
Mast cells
Megakaryocytes
Melanocytes
Cell Type Proportion
Time after injury in each tissue
Palate
Skin
Time
0 hours
6 hours
Day 1
Day 3
Day 7

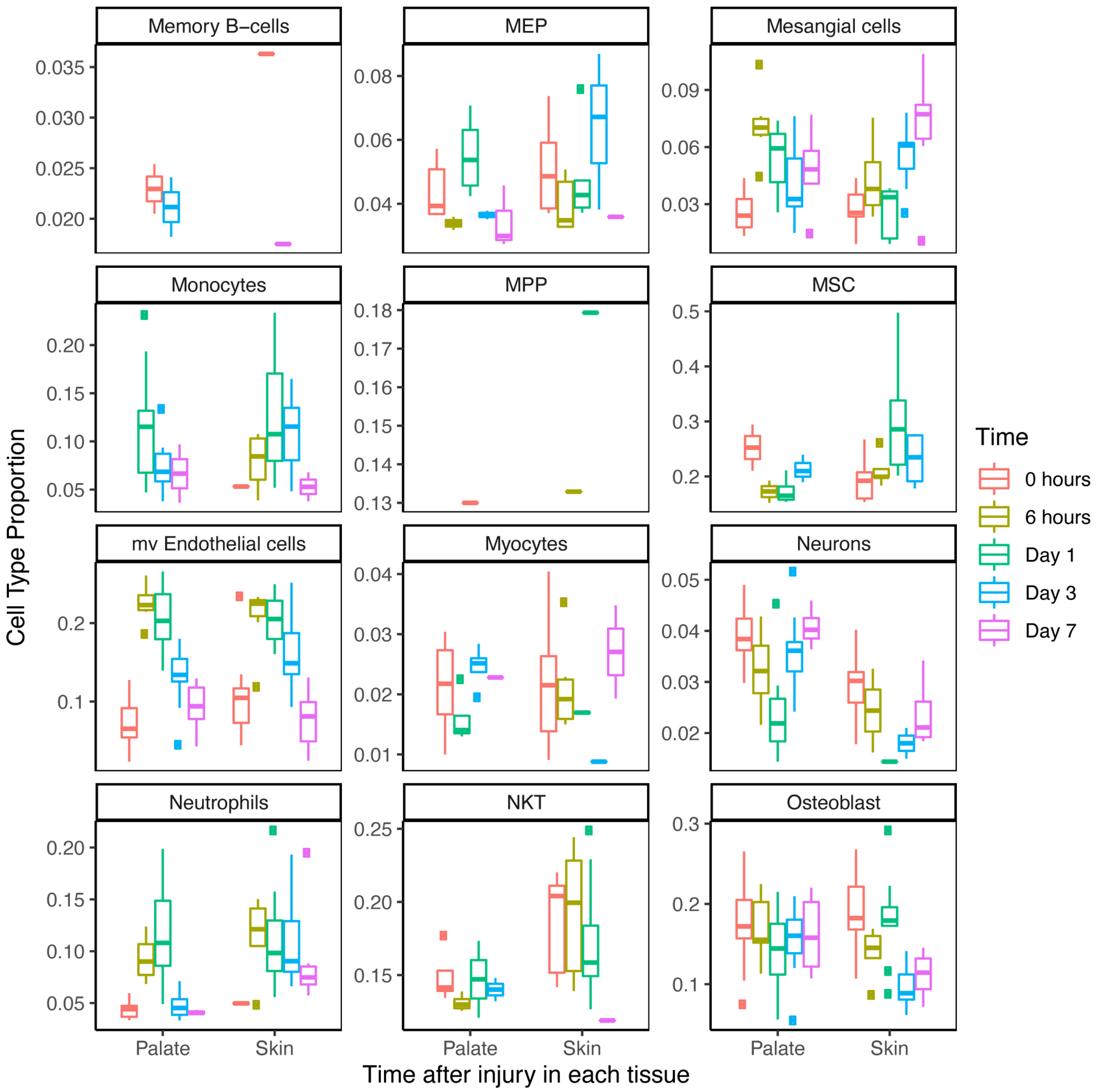
Memory B–cells
MEP
Mesangial cells
Monocytes
MPP
MSC
mv Endothelial cells
Myocytes
Neurons
Neutrophils
NKT
Osteoblast
Cell Type Proportion
Time after injury in each tissue
Palate
Skin
Time
0 hours
6 hours
Day 1
Day 3
Day 7

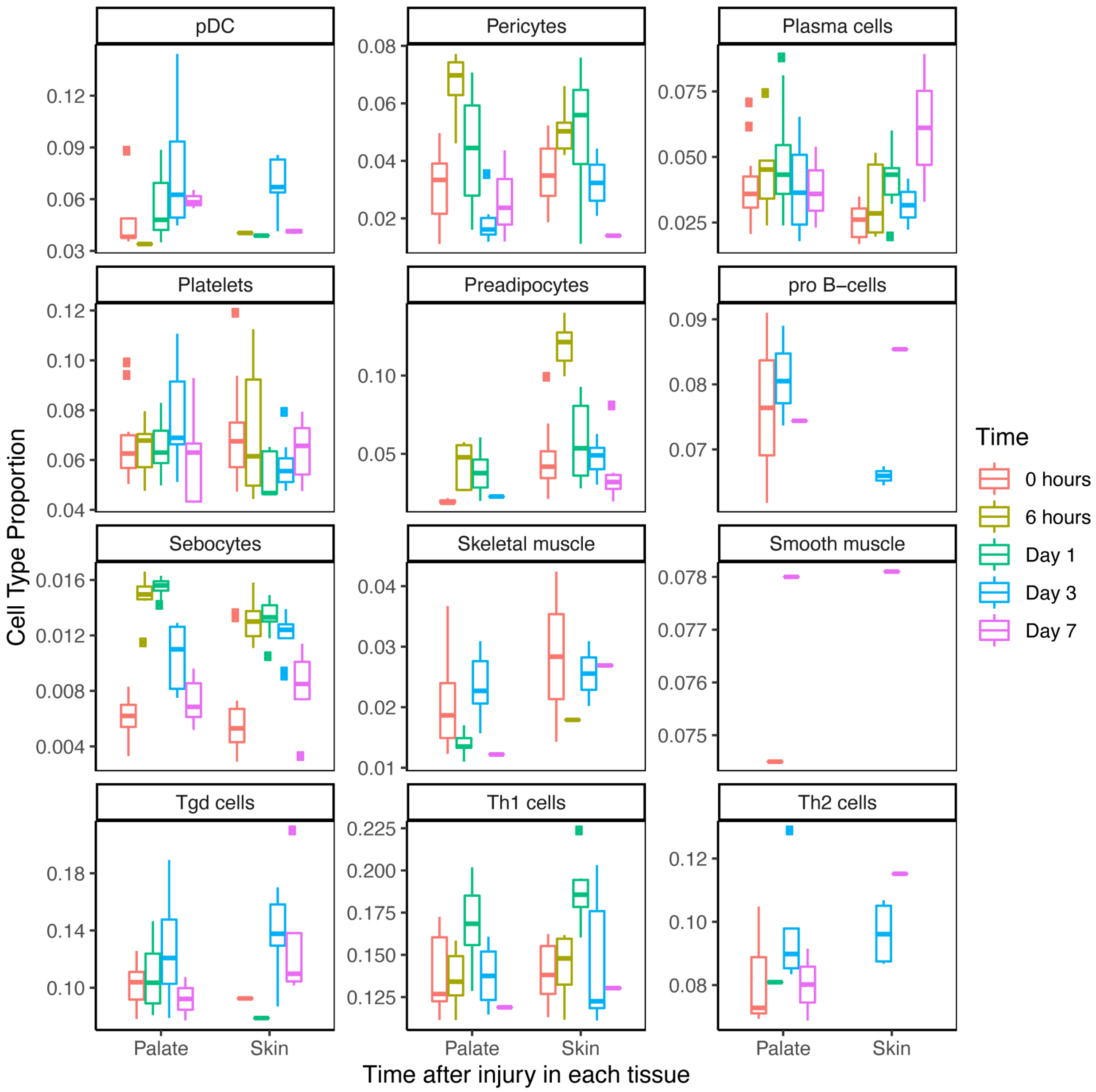
pDC
Pericytes
Plasma cells
Platelets
Preadipocytes
pro B−cells
Sebocytes
Skeletal muscle
Smooth muscle
Tgd cells
Th1 cells
Th2 cells
Cell Type Proportion
Time after injury in each tissue
Palate
Skin
Time
0 hours
6 hours
Day 1
Day 3
Day 7

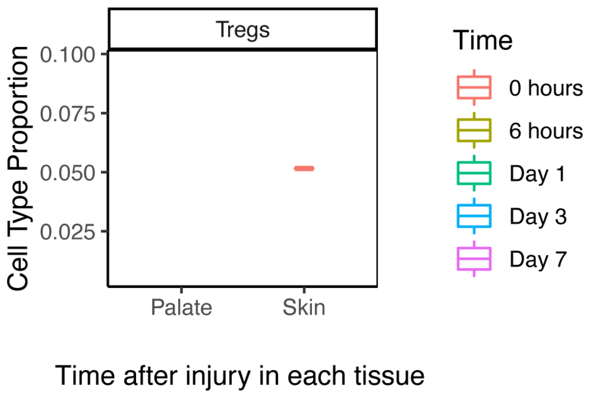


**Supplemental Figure 2. Cell deconvolution analysis of all 64 cell type proportions in tissue injury**

Results from the xCell deconvolution analysis were first filtered for statistical significance (p-value ≤ 0.2). Boxplots representing the cell type proportion (y-axis) at each point in time after injury in palate or skin (x-axis) were generated for each cell type (denoted by panel title). Colors represent the time point after injury, with 0 representing unwounded tissue. Outlier samples are represented by circles on the plots. Boxplots that do not appear for a specific tissue and time point indicate that the results from the xCell analysis did not reach statistical significance. All 64 cell types are included: (aDC = activated dendritic cells, CD4+ Tcm = CD4+ central memory T-cells, CLP = common lymphoid progenitors, CMP = common myeloid progenitors, DC = dendritic cells, Tgd = gamma delta T-cells, GMP = granulocyte-macrophage progenitors, HSC = hematopoietic stem cells, iDC = immature dendritic cells, ly endothelial cells = lymphatic endothelial cells, MSC = mesenchymal stem cells, mv Endothelial cells = microvascular endothelial cells, MPP = multipotent progenitors, NKT = natural killer T-cells, pDC = plasmacytoid dendritic cells, Tregs = regulatory T-cells, Th1 cells = Type 1 T-helper cells, Th2 cells = Type 2 T-helper cells, cDC = conventional dendritic cells).
